# R2DT: a comprehensive platform for visualising RNA secondary structure

**DOI:** 10.1101/2024.09.29.611006

**Authors:** Holly McCann, Caeden D. Meade, Loren Dean Williams, Anton S. Petrov, Philip Z. Johnson, Anne E. Simon, David Hoksza, Eric P. Nawrocki, Patricia P. Chan, Todd M. Lowe, Carlos Eduardo Ribas, Blake A. Sweeney, Fábio Madeira, Stephen Anyango, Sri Devan Appasamy, Mandar Deshpande, Mihaly Varadi, Sameer Velankar, Craig L. Zirbel, Aleksei Naiden, Fabrice Jossinet, Anton I. Petrov

## Abstract

RNA secondary (2D) structure visualisation is an essential tool for understanding RNA function. R2DT is a software package designed to visualise RNA 2D structures in consistent, recognisable, and reproducible layouts. The latest release, R2DT 2.0, introduces multiple significant features, including the ability to display position-specific information, such as single nucleotide polymorphisms (SNPs) or SHAPE reactivities. It also offers a new template-free mode allowing visualisation of RNAs without pre-existing templates, alongside a constrained folding mode and support for animated visualisations. Users can interactively modify R2DT diagrams, either manually or using natural language prompts, to generate new templates or create publication-quality images. Additionally, R2DT features faster performance, an expanded template library, and a growing collection of compatible tools and utilities. Already integrated into multiple biological databases, R2DT has evolved into a comprehensive platform for RNA 2D visualisation, accessible at https://r2dt.bio.

## INTRODUCTION

RNA is a fundamental molecule involved in various biological processes, such as translation, splicing, and immune response. The versatile nature of RNA is attributed to its ability to adopt complex tertiary (3D) structures, which facilitate its diverse functions. However, the intricate 3D structures make it challenging for non-specialists to examine RNA structure. Therefore, 2D diagrams are widely used to capture base pairing information and serve as a proxy for the complete RNA structure.

For many RNAs, it is essential to use standard, community-accepted orientations to ensure consistency and clarity in communication. For instance, transfer RNAs (tRNAs) are visualised in a cloverleaf shape with the 5′ end at the top left, while ribosomal RNAs (rRNAs) are commonly viewed using standard layouts introduced by the Gutell group (1). The usefulness of standardised layouts is not limited to well-studied RNAs, as it is often helpful to visualise related RNAs in similar orientations to facilitate structure comparisons (2).

While there are many software packages and websites for RNA 2D visualisation, including FoRNA (3), R2R (4), R2easyR (5), RiboDraw (6), RNArtist (7), RNAcanvas (8), RNApuzzler (9), RNAscape (10), VARNA (11), and others, these tools do not take advantage of the manually curated 2D diagrams familiar to users, and the resulting 2D structures are not guaranteed to have similar layouts for related sequences. R2DT is the only software that displays RNA structures in consistent, reproducible, and recognisable layouts and provides a comprehensive set of tools for creating, editing, and displaying RNA 2D diagrams (12).

R2DT, which stands for RNA 2D Templates, includes a comprehensive library of templates that represent a diverse range of RNA types and organisms (12). Using RNA sequence as input, R2DT searches the template library to choose the most suitable template and predicts a 2D structure compatible with it. The sequence and its predicted structure are then visualised using the Traveler software (13) following the layout of the chosen template.

This paper introduces R2DT 2.0, a new version of the software that incorporates multiple new features and improvements based on user feedback. The updated template library includes almost a thousand new templates and supports the RNArtist layout engine that minimises structural overlaps. The output diagrams can now be edited using specialised online software, including by using natural language prompts, and a new template-free mode makes it possible to visualise any given 2D structure. R2DT is integrated into multiple widely-used databases and is available as a web server, an API (14), a standalone program, and an embeddable web component, creating a comprehensive platform for RNA 2D structure visualisation.

## MATERIAL AND METHODS

### Overview of the R2DT pipeline

R2DT accepts input sequences in FASTA format, with an optional line for 2D structure in dot-bracket notation that can include pseudoknots. Originally developed for version 1.0, the general architecture of the R2DT pipeline involves two main steps: template selection and the generation of 2D diagrams.

#### Template selection

R2DT includes a library of 2D templates, each containing an RNA sequence, its 2D structure, and x, y coordinates for every nucleotide. If no 2D structure is provided by the user, then the input sequences are searched against the template library in several stages. Any sequences that do not find a match at any given stage are passed on to the following stages, with a sequence matching no more than one template.

First, each input sequence is compared to the larger RNA templates (currently rRNA and RNase P) using BLAST (15). Then each sequence is compared against profile HMMs of a subset of Rfam families using Ribovore (26). Finally, the tRNA models are searched using tRNAScan-SE 2.0 (16) which internally uses Infernal searches against a panel of tRNA covariance models and includes heuristic for pseudogene detection. An additional tRNA search with the Rfam tRNA model (RF00005) is also used to capture any tRNA sequences that have not been annotated by tRNAScan-SE. For sequences matching templates, the 2D structure is predicted using the Infernal `cmalign` program by aligning the input sequences to the template covariance models.

If a 2D structure is provided by the user, a new template is dynamically created using R2R (4) or RNArtist (7) (see below for more information about the new template-free mode). Template selection can also be bypassed by specifying the template directly via the command line or in the web interface.

#### Diagram creation

The 2D diagrams are created using the Traveler software, which uses the Infernal alignment (if available) to map input sequence to the template coordinates. Traveler computes the 2D structure layout, dynamically rearranging the structure to accommodate any insertions or deletions (12).

In addition to the standard 2D diagrams, a simplified representation of the structure is also produced, with a continuous line connecting each nucleotide in the sequence. These simplified diagrams are useful as thumbnail images for web pages or schematic diagrams.

#### Pipeline implementation

The R2DT pipeline is implemented in Python and is containerised using Docker to facilitate dependency management and ensure reproducibility across platforms. The Docker images are built using GitHub Actions and pushed to Docker Hub, with different R2DT versions available under stable tags. The R2DT Docker images are compatible with Singularity, LSF, Slurm, Kubernetes, and other computing environments, and have been successfully deployed in production within the EMBL-EBI’s Job Dispatcher framework (14) that powers the R2DT API.

A comprehensive test suite was developed to prevent breaking changes. The tests are executed as part of the pipeline and involve comparing previously generated images with newly produced ones. The comparisons are performed by converting the images to grayscale, computing per-channel differences, and using the Structural Similarity Index (SSI). Minor differences, below a predefined threshold, are accepted as normal, such as changes in font size or slight shifts in image elements. However, if code changes result in a significantly different 2D diagram, the test suite fails. This process ensures that regressions are detected automatically and early in the development cycle.

#### Template library

R2DT includes a template library with 4,612 2D templates. An Infernal covariance model (17) is built for each template using its sequence and 2D structure. These models are used for both matching input sequences to templates and for predicting the 2D structure compatible with the selected template. The template library is continuously updated as new Rfam (18) releases become available or new templates are submitted by the community. The current composition of the template library is discussed in more detail in the Results section.

## RESULTS

### New types of visualisations

#### Visualising annotations as data layers

RNA 2D structures are often used to present experimental results, such as SNPs, sequence conservation, or reactivity. The new version of R2DT enables users to visualise any position-specific information as data layers. Each nucleotide of the input sequence can be annotated with numeric or textual labels, and a mapping between colours and labels can be provided.

Internally R2DT uses this functionality to visualise the alignment quality between a query sequence and a template. The quality is represented by posterior probabilities calculated by the Infernal software (17). Nucleotides with a high probability of alignment (95-100%) are not highlighted while the rest of the nucleotides are coloured according to the alignment confidence using a colour-blind safe palette (19).

An example diagram annotated with posterior probabilities is shown in Figure 1A, where the coloured circles help to pinpoint species-specific differences between the human and rabbit ribosomes. Figure 1B displays DMS reactivities for stem loops (SL) 1 through 4 from SARS-CoV-2 (20), with higher reactivities corresponding to the nucleotides not involved in Watson-Crick base pairs.

**Figure 1:**
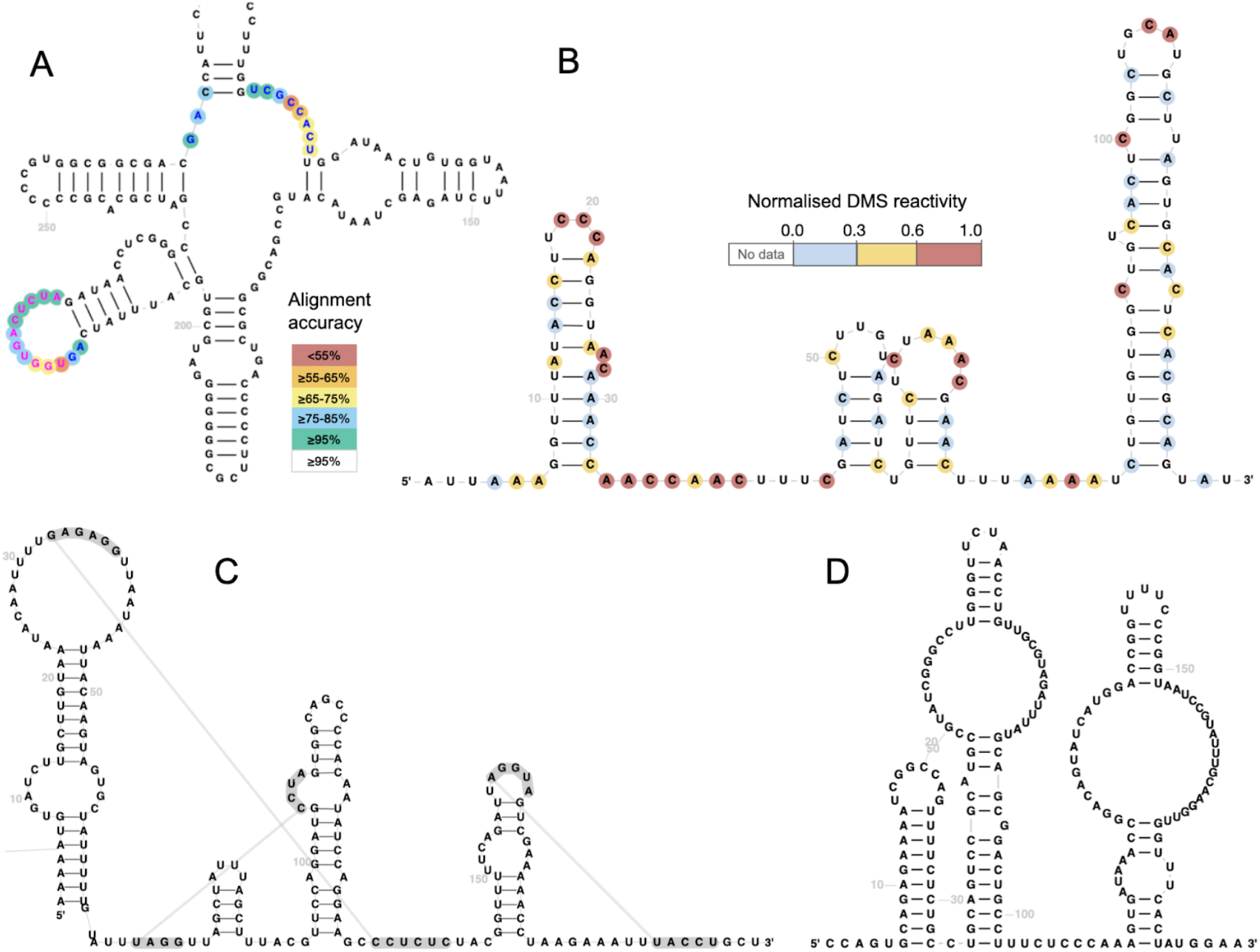
A) A fragment of a rabbit small ribosomal subunit rRNA sequence visualised using the human template with Infernal posterior probabilities reflecting alignment confidence shown as coloured circles (RNAcentral ID [URS000086855E_9986](https://rnacentral.org/rna/URS000086855E/9986)). The highlighted regions help identify species-specific differences between the human and the rabbit rRNAs. B) Normalised DMS reactivities for stem loops SL1-SL4 of SARS-CoV-2 (data from (20)). C) Cripavirus IRES with three pseudoknots (RNAcentral ID URS0000A7638F_12136). D) Bridge RNA from *Escherichia coli* (INSDC accession CP147105.1/771271-771455) visualised using the new template-free mode (data from (21)).

#### Pseudoknot visualisation

RNA pseudoknots are structural motifs where non-nested base pairing interactions form between non-contiguous regions of the sequence, creating a knot-like structure that plays critical roles in RNA folding and function. The original version of R2DT was not able to display pseudoknots, but an updated version of the Traveler software enabled R2DT to include them in the diagrams. The pseudoknots annotations are either provided by users or are extracted from the Rfam consensus 2D structures. Upon alignment of a target sequence to the covariance model using Infernal, the pseudoknot annotation is transferred to the alignment using the `cmalign` program and included in the final 2D diagram. Any number of pseudoknots can be displayed, for example, Figure 1C shows a Cripavirus IRES with three pseudoknots.

#### Template-free mode

While the key strength of R2DT is the ability to reproduce a manually curated template, it is also necessary to be able to visualise RNA structure without pre-defined templates. R2DT 2.0 has a new template-free mode that accepts a sequence and a 2D structure in dot-bracket notation. The structure is visualised using the R2R (4) or RNArtist (7) software, and the resulting diagram is used to generate a template and a 2D diagram. Figure 1D shows a bridge RNA from E.coli that was recently discovered (21) and does not yet have a template, but can still be visualised in a template-free mode.

The output of the template-free mode follows the standard R2DT format, so it can be used to display position-specific information or edited by all software compatible with R2DT (see below). The template can be used to visualise other sequences in the same layout, which can be useful for displaying sequence search results or comparing related RNA structures. Users can submit new templates to the R2DT template library using GitHub or build their own template libraries for applications requiring privacy (see the documentation for detailed instructions). The template-free mode has already been successfully used to visualise RNA structures found in the HCV genome (22) and in SHAPEwarp-web (23) to visualise SHAPE reactivities.

The template-free mode significantly expands the utility of R2DT, as it can be used to visualise any 2D structure, similar to programs like VARNA (11) or FoRNA (3), but with added support for interactive editing and other R2DT-specific features described below.

### Interactive editors for R2DT diagrams

In previous versions, users often requested the ability to modify R2DT output to adjust layouts or add custom annotations. While R2DT produces Scalable Vector Graphics (SVG) files that can be edited using any software that supports the SVG file format, such as Adobe Illustrator or GIMP, these tools lack specialised features needed for RNA-specific tasks. To address this, two web-based RNA structure editors, RNAcanvas (8) and XRNA-React, were integrated with R2DT to facilitate the creation of publication-quality images (Figure 2). Both editors enable users to import/export 2D structures in multiple formats and select, edit, and format the selected 2D layout and base pairs, offering a range of features for comprehensive RNA editing and visualisation. Notably, the editors can be used to create new R2DT templates (see the RNA 2D JSON Schema format section).

**Figure 2:**
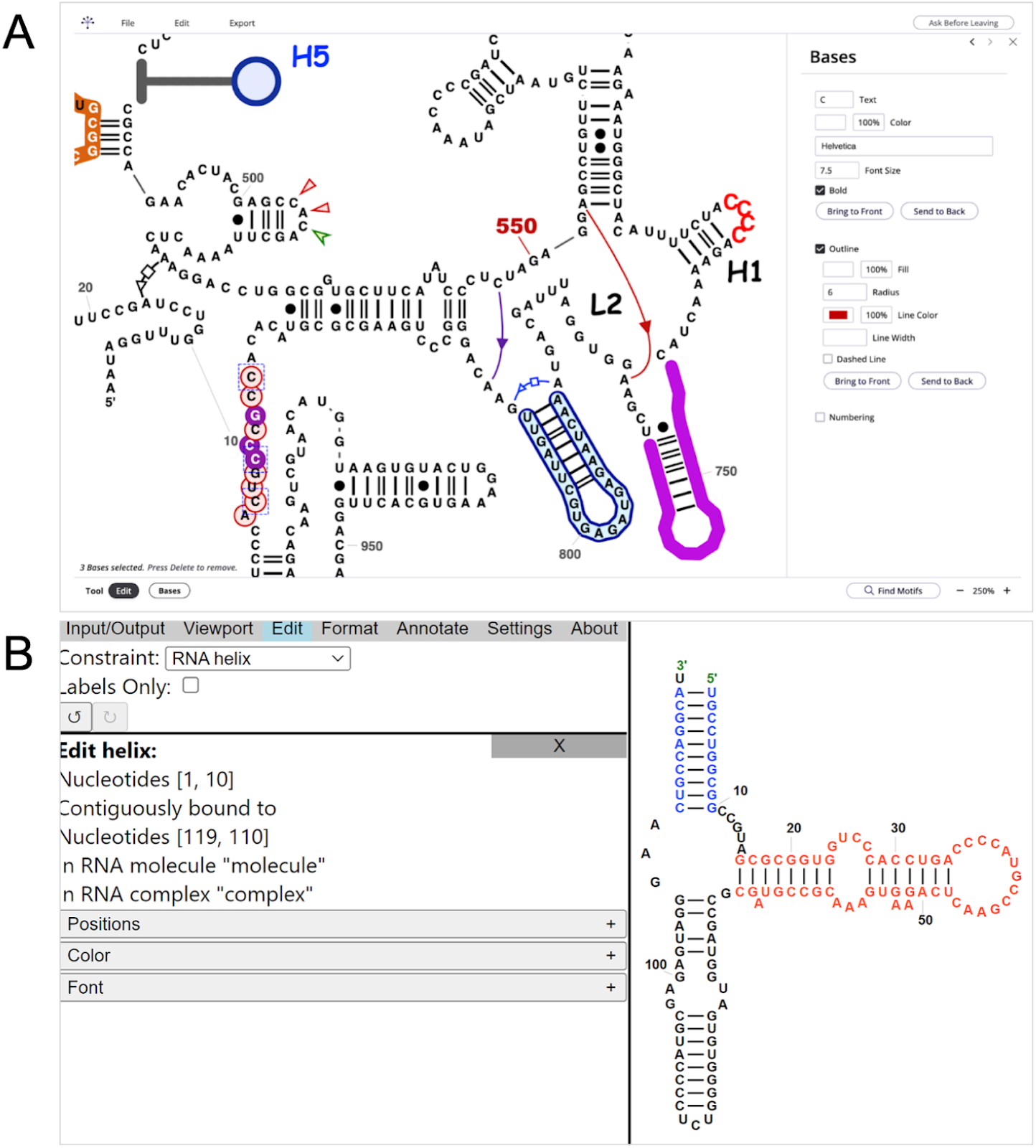
A) Screenshot of the RNAcanvas user interface showing human mitochondrially encoded 12S rRNA (MT-RNR1). The layout and the 2D structure were generated by R2DT, while the fonts, colour, background, and non-canonical interactions were configured in RNAcanvas. B) Screenshot of the XRNA-GT website showing a 5S rRNA.

#### XRNA-React

XRNA-React, available at https://ldwlab.github.io/XRNA-React, is a web-based tool for editing and visualising RNA secondary structures. It is an expanded version of the original XRNA software developed by Harry Noller (available at http://rna.ucsc.edu/rnacenter/xrna/xrna.html but compatible only with a deprecated version of Java). The tool provides several editing modes, such as selecting single nucleotide, single strand, base pair, helix, RNA sub-domain, and entire RNA molecule, allowing users to make precise adjustments to the structure.

#### RNAcanvas

RNAcanvas (8) is available at https://rnacanvas.app. Similar to XRNA-React, it enables rich editing functionality and provides a high level of customisation for fonts, background colours, labels, and other secondary structure elements. RNAcanvas has a built-in motif search tool that facilitates quick identification of sequences of interest.

#### RNA 2D JSON Schema format

The interactive editors leverage a new format called RNA 2D JSON Schema that provides a standardised way of storing one or more RNA sequences, 2D structure, base pairs in the Leontis-Westhof notation (24), x, y coordinates, as well as styling metadata. The file format is implemented using JSON Schema (https://json-schema.org/), a widely adopted standard for data exchange that supports validation and consistency checks.

R2DT generates RNA 2D JSON Schema files as part of its standard output, and both RNAcanvas and XRNA-React can import RNA 2D JSON Schema files from local files or via URLs (for example, the R2DT API or any public file). The modified diagrams can then be exported in RNA 2D JSON Schema format.

Conversely, R2DT can use RNA 2D JSON Schema files as input in order to generate new templates, enabling new types of workflows. Users can visualise 2D structure using R2DT in template-free mode, interactively edit the resulting diagram using RNAcanvas or XRNA-React, download the results as RNA 2D JSON Schema, and create new R2DT templates for private use or submission to the R2DT library (see https://docs.r2dt.bio/en/latest/templates.html for more information).

We encourage the adoption of this format to foster the development of a robust ecosystem of tools for producing and editing RNA 2D structure diagrams. The schema description is freely available on GitHub (https://github.com/LDWLab/RNA2D-data-schema/) and can be used by any software that generates RNA 2D structures.

#### Using natural language for editing RNA 2D diagrams

The rise of large language model (LLM) tools, such as ChatGPT, capable of code generation in response to natural language inputs, streamlines the creation of natural language interfaces to programming applications, including RNA 2D structure software.

A separate version of RNAcanvas (termed RNAcanvas Code, available at https://code.rnacanvas.app) comes paired with a custom GPT that serves as an “AI assistant” and is capable of generating code that leverages the RNAcanvas Code API in response to natural language inputs from the user. This custom GPT, available at https://chatgpt.com/g/g-jh8gXtvrC-rnacanvas-ai-assistant (requires an OpenAI account to access), was created by inputting the documentation for the RNAcanvas Code API to the custom GPT.

RNAcanvas Code allows the user to interact with and edit RNA structure drawings by entering code (self-or LLM-generated) into the web browser console (a standard feature of all major web browsers). The users can ask the RNAcanvas custom GPT to load R2DT results in RNA 2D JSON Schema format or start with any other RNA 2D structure. Then the users can edit the structure, for example, by asking “Write code to colour all U residues red.” The user would copy and paste the code generated by the custom GPT into the web browser console of the RNAcanvas Code app to see the result and manually adjust the diagram, if needed.

Natural language queries from the user can be more complex, such as “Write code to find all instances of the motif “AGU”, colour them blue and increase their font size to 16 while maintaining the centre points of all bases.” Example prompts, their outputs, and the corresponding diagrams can be seen in Supplementary Information.

Although LLM-generated code is often useful, it can also be incorrect at this early stage of development and requires that the user be discerning when making use of it. As the LLM technology and the available tools continue to evolve, we anticipate that tools like RNAcanvas Code will be built directly into the user interface to simplify complex and repetitive tasks, without requiring users to interact with the underlying code.

### Pipeline improvements

#### Faster template selection using BLAST

In previous versions of R2DT, searching through thousands of templates created a performance bottleneck and increased the tool’s carbon footprint (25). The template selection process relied on the Ribovore software (26) which used profile hidden Markov model (HMM) searches. These searches were computationally intensive, especially when applied to large models like those for ribosomal RNAs.

In R2DT 2.0, a new BLAST-based (15) filtering step has been developed, significantly improving performance. For example, selecting a ribosomal template for a set of 3,708 distinct RNA sequences from the PDB (27) takes an average of 0.4 seconds using the new BLAST-based approach compared to 316.3 seconds with the previous method (tests conducted on an M2 MacBook Air with 24 GB of RAM). The BLAST-based method identified 816 templates whereas HMMs found 802.

While the templates selected by each method are not always identical, they are generally similar, and manual inspection of the diagrams did not reveal any significant issues. The Rfam models and tRNAs continue to be analysed with Ribovore and tRNAScan-SE 2.0 (16), respectively, as these templates typically derive from shorter RNAs, where BLAST searches may produce false negatives.

#### Constrained folding of regions that do not align to templates

While 2D templates are effective at providing the scaffold for 2D layouts, some sequences contain insertions that do not align to the templates, for example, species-specific extensions that are absent from the family consensus structure. By default R2DT displays such regions as unfolded loops (Figure 3A). In order to predict base pairs for the unfolded regions, R2DT can now use RNAfold from the Vienna RNA package (28). There are three ways of using constrained folding:

**Figure 3.**
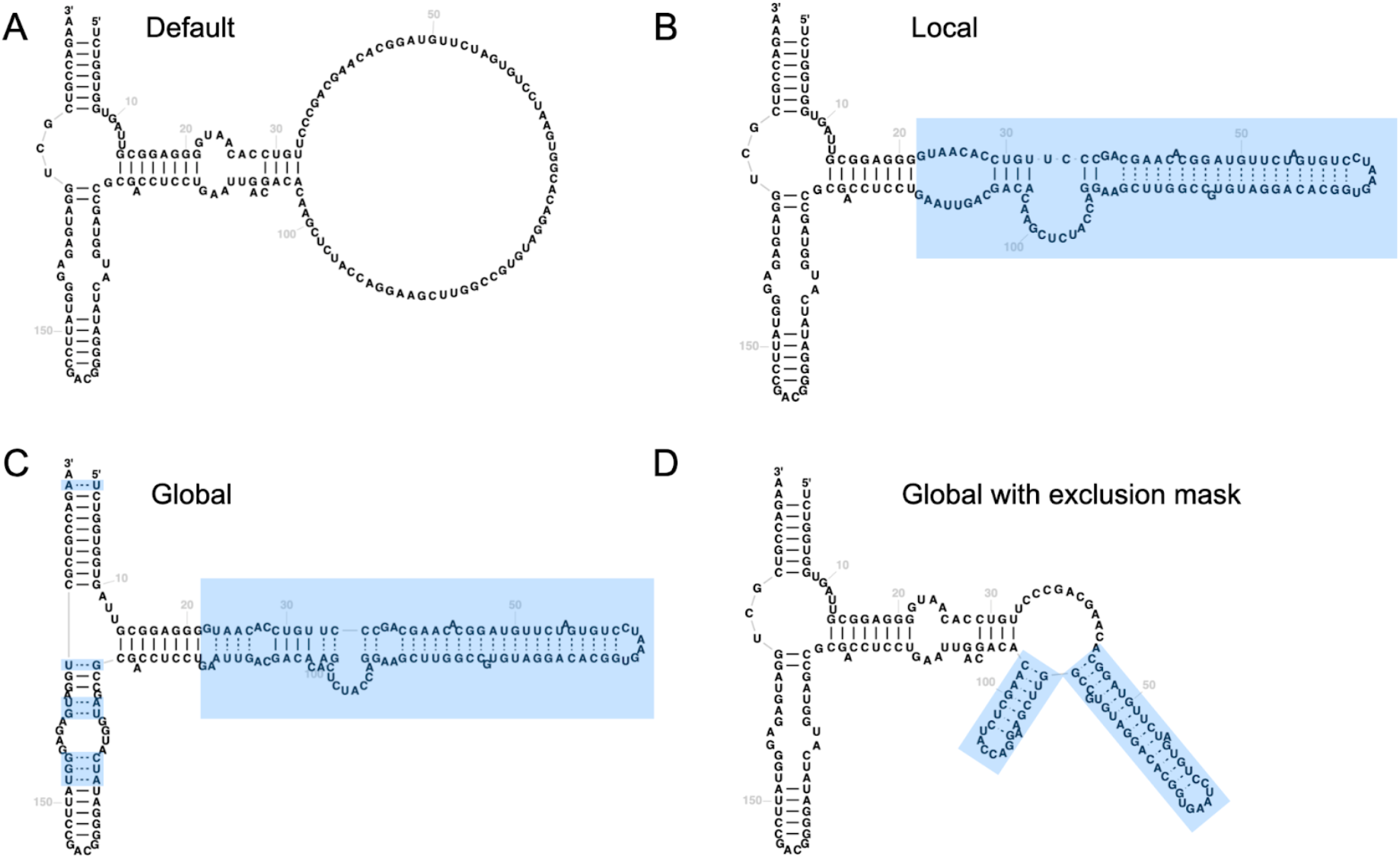
Example diagrams illustrating different constrained folding modes in R2DT. The 5S sequence from *Halobacteroides halobius* is visualised using a 5S template from *Bacillus subtilis* (INSDC accession CP003359.1/20186-20360). A) Default R2DT output without constrained folding. B) Local folding mode where RNAfold is used to predict the structure for the insertion relative to the template. C) Global folding mode with additional base pairs predicted throughout the structure. D) An example of using global folding mode with an exclusion mask. In this case all unpaired nucleotides aligned to the template were prevented from forming base pairs. Predicted base pairs are shown as dashed lines. Changes relative to default output are highlighted in blue boxes.

1. Local folding: the 2D structure of the insertion relative to the template is predicted with RNAfold and added to the diagram (Figure 3B).
2. Global folding: the entire molecule is folded using RNAfold with the template structure provided as a constraint (Figure 3C).
3. Global folding with masking: the entire molecule is folded using RNAfold, but the user can provide an arbitrary exclusion mask to prevent any specific nucleotide from forming base pairs. For example, in Figure 3D all nucleotides matching the template preserve their original base pairing state as specified in the template, so that the nucleotides that were unpaired stay unpaired.

### New and updated templates

To improve the quality of the output diagrams and keep up with the new RNAs being discovered, the R2DT template library continues to grow. Compared to 3,647 templates in the first version, the number of templates increased by 965 in R2DT 2.0, with Rfam being the largest source of new templates.

For example, three new templates were developed for the archaeal Type T RNase P RNA representing the variants in different phylogenetic clades, namely *Caldivirga, Pyrobaculum/ Thermoproteus*, and *Vulcanisaeta* (29). These templates replace the Rfam-based template from family RF02357. In addition, new rRNA templates were included for tomato (*Solanum lycopersicum*) (30), *Leishmania donovani* (31) and *Trypanasoma brucei* (32, 33).

While the majority of templates (80%) are generated automatically, manual curation continues to be essential for large diagrams and RNAs that are commonly visualised in standard orientations, such as rRNA and tRNA. All templates for R2DT 2.0 are summarised in Table 1.

**Table 1.** The RNA 2D structure template library in R2DT 2.0.

| RNA type | Template source | Number of templates | Manually curated? |
| --- | --- | --- | --- |
| Small RNAs | Rfam (18) | 3,805 | No |
| rRNA | CRW (1) | 661 | Yes |
|  | RiboVision (34) | 31 | Yes |
| tRNA | GtRNAdb (16, 35) | 91 | Yes |
| RNase P | RNase P database (29, 36, 37) | 24 | Yes |
|  | Total | 4,612 |  |

#### RNArtist templates for Rfam families

R2DT automatically generates templates for thousands of Rfam families using R2R, but these templates can sometimes have overlapping structural elements (Figure 4, left subpanels). To overcome this limitation, R2DT now supports RNArtist, a software package designed to minimise overlaps. RNArtist templates are precomputed for all Rfam families, and R2DT automatically chooses between the R2R and RNArtist layouts to minimise the number of overlaps (Figure 4, right subpanels). Users can also specify their preferred layout via a command line option. The RNArtist software can also be used to generate new layouts in the template-free mode (Figure 1D).

**Figure 4.**
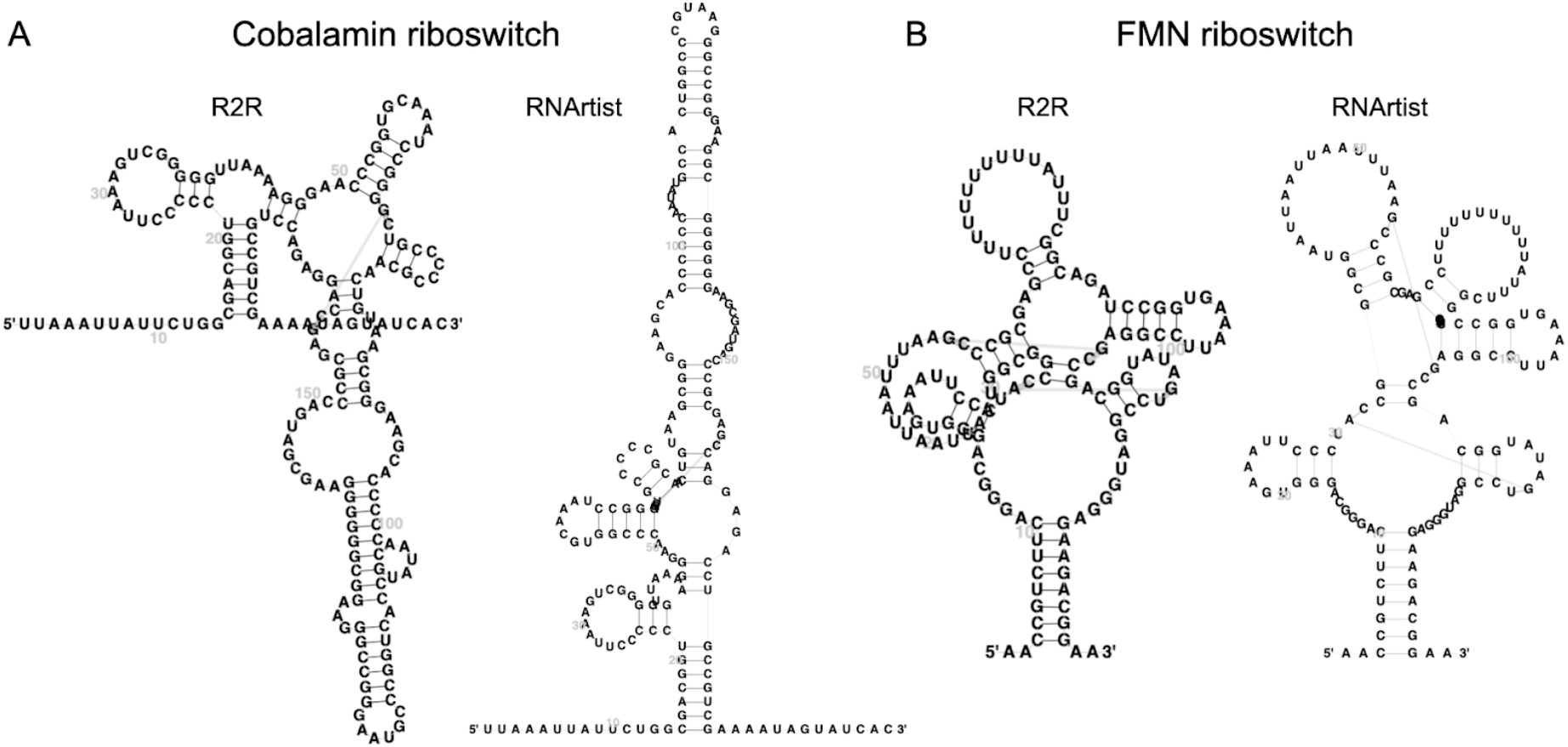
Example R2DT templates based on consensus 2D structures of Rfam families visualised using R2R and RNArtist in the left and right subpanels, respectively. A) Cobalamin riboswitch (Rfam ID RF00174). B) FMN riboswitch (Rfam ID RF00050). The R2R layouts contain major overlaps between the structural elements making it difficult to see the nucleotides, while the RNArtist layouts prevent the overlaps and are more legible. For both families, R2DT 2.0 uses the RNArtist layouts by default.

#### New tRNA templates

R2DT 2.0 has also made notable improvements to the isotype-specific tRNA templates based on the tRNAscan-SE 2.0 models (16). These templates include slight adjustments to the consensus sequences and/or 2D structures of both the domain-specific and archaeal isotype-specific tRNA templates. Additionally, each residue within the templates includes a numbering-label field that represents the commonly used Sprinzl positions for tRNAs (38). The numbering labels can be propagated from the template to the resulting structure using a new option in the Traveler software. However, as certain positions may or may not exist depending on the isotype and phylogenetic clade, the numbering labels are not displayed at that region using the new Traveler options (`-l --numbering “13,26”`).

A new set of 23 templates has been created for mitochondrial tRNAs in vertebrates. These templates correspond to the covariance models named as TRNAinf-mito-vert-*.cm or the CM database TRNAinf-mito-vert, which are included in tRNAscan-SE starting with v2.0.10.

Unlike cytosolic tRNAs, which have one template per isotype, mitochondrial tRNAs have two templates for mt-tRNA-Leu and mt-tRNA-Ser, separated by anticodons. Furthermore, mt-tRNA-Cys has two templates/CMs, one with a typical cloverleaf structure and the other lacking a D-arm (found in armadillo and some reptiles). These improvements to the isotype-specific tRNA templates in R2DT 2.0 and the creation of new templates for mitochondrial tRNAs in vertebrates will enhance the accuracy and efficiency of tRNA visualisation.

### Overview of resources integrating R2DT

The R2DT visualisations have been successfully incorporated into multiple online resources, collectively reaching tens of thousands of users each month. For example, the RNAcentral database displays precomputed diagrams for over 25 million ncRNA sequences (39), and R2DT structure diagrams are automatically generated for RNAcentral sequence search results in RNAcentral and several other websites relying on RNAcentral sequence search, such as Rfam (18), GtRNAdb (35), and snoDB (40). The `r2dt-web` and `pdb-rna-viewer` web components (see below) facilitate embedding R2DT diagrams into any website, enabling FlyBase (41), PomBase (42), SGD (43), RiboVision2 (34), and NAKB (44) to present detailed 2D structure diagrams on their pages. Several standalone software packages also integrate with R2DT, including SHAPEwarp-web (23) and RNAvigate (45) that show SHAPE activities overlaid on the 2D structure diagrams.

#### Case study: Displaying RNA 2D structures in PDBe

A special visualisation based on R2DT has been developed to enable seamless integration between RNA sequence, 2D and 3D structures from the PDB (27) on the Protein Data Bank in Europe (PDBe) web site (46). Base pairing annotations derived from annotating the 3D structures with FR3D Python (47) are superimposed on the R2DT-generated layouts, using the Leontis-Westhof base-pairing classification (24). Standard Watson-Crick base pairs appear in the viewer as lines connecting the nucleotides (Figure 5A, left). In contrast, non-Watson-Crick base pairs are depicted using Leontis-Westhof notation to emphasise various interaction types within the 2D structure elements (Figure 5B, left). Clicking on bases in the 2D diagram directs users to the corresponding nucleotides in the 3D molecular graphics viewer, Mol* (Figure 5, right panels) (48). Further interaction with the specific base in Mol* reveals the local region in greater detail, including the base pairing interactions between nucleic acid bases.

**Figure 5.**
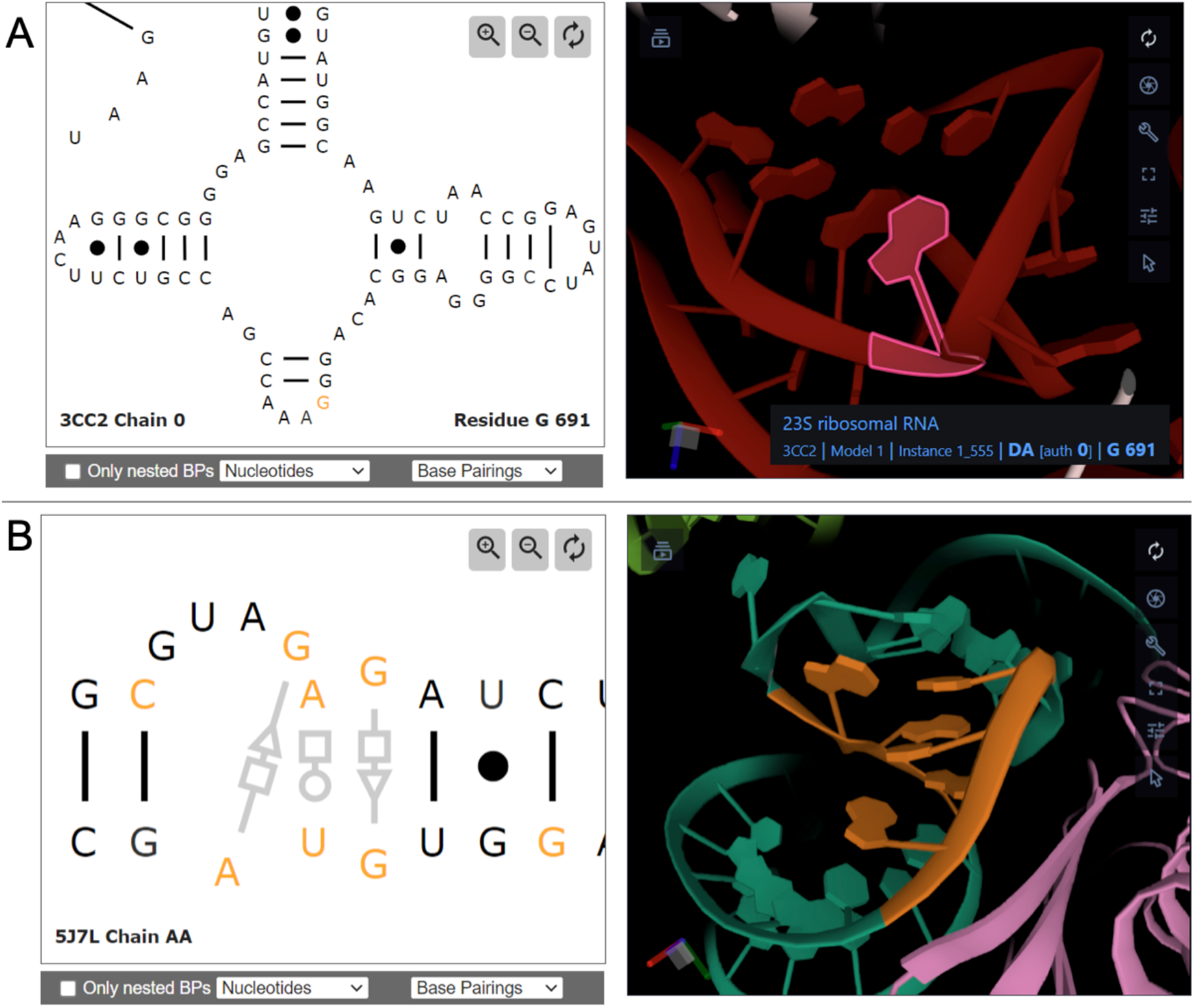
A) Screenshot of the 2D and 3D components on the macromolecule page for 23S ribosomal RNA from *E. coli*, in PDB entry 3CC2. The nucleotide G691 is selected in 3D and also highlighted with orange in 2D. B) Screenshot of the 2D and 3D components on the macromolecule page for 16S ribosomal RNA from *E. coli*, in PDB entry 5J7L, highlighting a kink-turn RNA 3D module and its non-canonical base pairs annotated using the Leontis-Westhof nomenclature (24).

### R2DT utilities

#### Reusable web components

R2DT is integrated into several reusable web components that can be used to view and edit its output. For example, the `pdb-rna-viewer` widget powers the PDBe interface. The data for the web component is available via static URLs for RNA chains in the PDB (base pair information and layout data). Both FR3D Python and R2DT have been integrated into the weekly release process of PDBe, ensuring that the data is available for every RNA 3D structure on the day of the weekly PDB release.

The `r2dt-web` component can be used to display 2D diagrams given an RNAcentral unique RNA sequence (URS) identifier. The precomputed structures and layouts are loaded from the RNAcentral database. RNAcanvas Code can also be embedded into any page and allows loading RNA 2D JSON Schema generated by R2DT, in addition to drawing RNA structures independent of R2DT using sequence and dot-bracket notation.

#### Animated visualisations showing structural rearrangements

The R2DT functionality can be extended by lightweight utility programs that make R2DT easier to use or apply it in new contexts. For example, an R2DT utility `animate.py` helps create animated SVG images that demonstrate alternative 2D structures of the same size, dynamically morphing one into another. While not intended to convey kinetic or topological information, such animations can facilitate the comparison of 2D structures, as well as display structural heterogeneity, riboswitches, or riboSNitches (49). Video 1 shows an example of animated transition between a reference and a solution structure from RNA Puzzle 39, showing which nucleotides from the predicted three-way junction form an additional helix in the reference four-way junction structure.

**Video 1**. The animation shows the 2D structure of the reference model from RNApuzzles Puzzle 39 and the 2D structure of a predicted model (Dfold group, model 3). The base pairs were extracted from the PDB files using the RNAView software.

#### Python R2DT API client

In some use cases it is desirable to generate RNA 2D structures without installing R2DT or running any computationally intensive tasks. The `r2dt-client` is a separate Python package that provides a convenient interface for R2DT web API and facilitates integration of RNA 2D structure diagrams into Jupyter notebooks, Google Colab, or any other environment that can execute Python code. While similar to the `draw_rna` tool from the Das lab (https://github.com/DasLab/draw_rna), `r2dt-client` is a wrapper that uses the R2DT API to compute the diagrams remotely while `draw_rna` runs locally. The package is released via PyPi and the documentation is available at https://github.com/r2dt-bio/r2dt-client.

## DISCUSSION

The new features introduced in R2DT 2.0 represent significant improvements compared with previous versions (12) and other software (3–10), addressing key limitations and expanding R2DT’s applicability to new use cases. The ability to display position-specific information makes R2DT a more versatile tool for researchers working with diverse RNA species, particularly in complex and data-rich contexts such as SNP mapping and SHAPE reactivity analysis. The introduction of the template-free mode removed the reliance on pre-existing templates, helping visualise novel or less-characterised RNAs. The successful application of this mode to the HCV genome and SHAPEwarp-web underscores its potential for broader adoption in RNA research.

An important outcome of the R2DT project is the creation of the RNA 2D JSON Schema format, which is used for storing and editing 2D structures. It provides a common framework that enables RNA visualisation software to be used interchangeably. It also allows one to use the interactive editors that can load RNA 2D JSON Schema files from any public URL. For example, by generating RNA 2D JSON Schema files, 2D structure prediction software could enable its users to edit the resulting diagrams in the same way as R2DT.

One of the primary strengths of R2DT is its extensive and ever-growing library of manually curated RNA templates, which are essential for producing accurate and consistent 2D structure diagrams. In future versions, continuous improvement efforts will focus not only on increasing the number of templates available but also on enhancing their quality. This includes refining existing templates to better reflect updated biological data (for example, new experimentally determined 3D structures), creating new templates for recently discovered or previously unrepresented RNA families, and integrating emerging sources of high quality 2D diagrams, such as Ribocentre-switch (50). These improvements will ensure that R2DT remains capable of handling the diverse range of RNA molecules being studied across different species and research contexts.

Automatically generating clear and legible diagrams remains a key area for improvement, and integrating additional layout engines, such as RNApuzzler (9), can help produce intersection-free drawings. Another challenge is to simplify R2DT installation without reliance on Docker containers, which can be a barrier for some users. Facilitating R2DT installation via conda, adopting cross-platform technologies such as the Electron framework, or enabling R2DT to run directly in web browsers through WASM technology could greatly enhance the installation experience.

Recognising the importance of interoperability, R2DT will continue to expand its integration with other tools and biological databases, aiming to create a cohesive ecosystem where RNA 2D structures can be easily visualised, edited, and analysed across different platforms. Maintaining consistent 2D representation across biological resources means that users do not have to learn new ways to view the same RNA molecule on different websites.

As RNA 3D structure prediction software methods mature and the experimental techniques such as cryo-EM become more widely used for RNA structure determination (51), we will work with 3D structure prediction software developers, the RNA Puzzles team (52), and PDBe to use R2DT for viewing 2D structure, in order to enable efficient exploration and comparison of reference and predicted structures. We also plan to integrate R2DT with genome browsers and other relevant data sources. This integration will enable users to overlay data from genome tracks, such as VCF or BigWig files to represent conservation, variation, and other annotations directly mapped onto RNA 2D structure diagrams.

We are also exploring the use of large language models (LLMs) for manipulating RNA 2D structure. By using simple textual commands rather than multiple manual steps, this approach aims to make the process more intuitive and reproducible, thereby lowering the learning curve for users unfamiliar with diagram-editing tools. With advancements in agentic AI, there is potential to develop systems where AI agents autonomously generate and update R2DT templates, maximising legibility and integrating the latest data.

As RNA continues to be a focal point in molecular biology, the ability to accurately and efficiently visualise RNA structures in a consistent way is crucial. R2DT 2.0 not only addresses the current needs but also lays the groundwork for future developments, ensuring that it remains a useful tool for the exploration and understanding of RNA function and structure. Overall, the enhancements in R2DT 2.0 and the development of compatible tools and utilities make it a robust and flexible platform for RNA 2D structure visualisation, with applications ranging from detailed structural annotation to large-scale comparative analyses. Its user-friendly features, combined with the ability to handle complex and novel RNA structures, position R2DT 2.0 as an essential resource for both experienced RNA researchers and those new to the field. We welcome community feedback and contributions at https://r2dt.bio.

## DATA AVAILABILITY

The R2DT web interface is available at https://r2dt.bio and https://rnacentral.org/r2dt. The R2DT source code is available at https://github.com/r2dt-bio/r2dt and precomputed Docker images are available at https://hub.docker.com/r/rnacentral/r2dt. The `r2dt-web` embeddable widget is available at https://github.com/rnacentral/r2dt-web. The `pdb-rna-viewer` is found at https://github.com/PDBeurope/pdb-rna-viewer. The `r2dt-client` package is available at https://github.com/r2dt-bio/r2dt-client. The RNA 2D JSON Schema is available at https://github.com/LDWLab/RNA2D-data-schema. The latest information can be found in the R2DT documentation at https://docs.r2dt.bio.

## SUPPLEMENTARY DATA

Supplementary Data are available at NAR online.

## ACKNOWLEDGEMENTS

The authors would like to thank the organisers and participants of the Computational Approaches to RNA Structure and Function meeting held at the Centro de Ciencias de Benasque Pedro Pascual, where many of the R2DT features were conceived, discussed, and implemented.

## VIDEOS

**Video 1.**
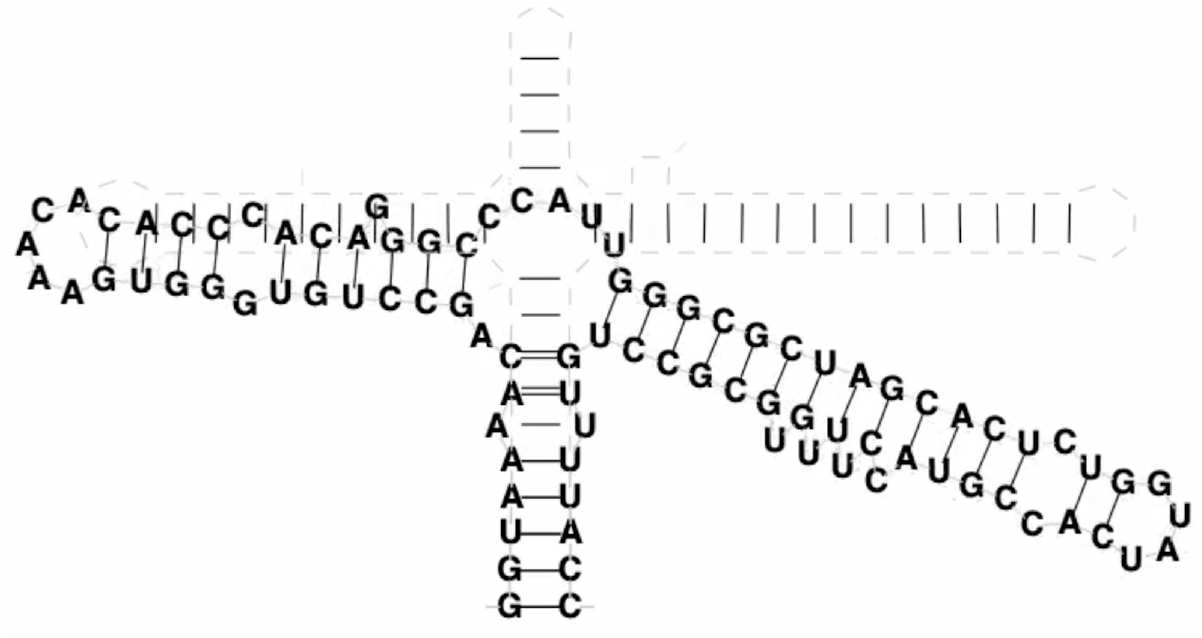
A still from the animation showing the 2D structure of the reference model from RNApuzzles Puzzle 39 and the 2D structure of a predicted model (Dfold group, model 3). The base pairs were extracted from the PDB files using the RNAView software. The video file is available in Supplementary Data.

