## Supplementary Information for "R2DT: a comprehensive platform for visualising RNA secondary structure"

### SUPPLEMENTARY DATA

#### Using R2DT and RNAcanvas Code for editing RNA 2D diagrams using natural language

The following steps demonstrate editing R2DT output in RNAcanvas Code using a custom ChatGPT assistant and natural language prompts.

**STEP 1.** Visualise the following bridge RNA secondary structure in R2DT at https://r2dt.bio:

```
>bridge_rna
CCAGUGCAGAGAAAAUCGGCCAGUUUUCUCUGCCUGCAGUCCGCAUGCCGUAUCGGGCCUUGGGUUCUAACCUGUUGCGUAGAUUUAUGCAGCGGACUGC
CUUUCUCCCAAAGUGAUAAACCGGACAGUAUCAUGGACCGGUUUUCCCGGUAAUCCGUAUUUGCAAGGUUGGUUUCACUAUGGAA
.....((((((((((........)))))))))).(((((((((.(((............
(((((....))))).............))))).)))))))))).......(.((((....(((.............
((((((.....))))))...)...............)))..)))).)....
```

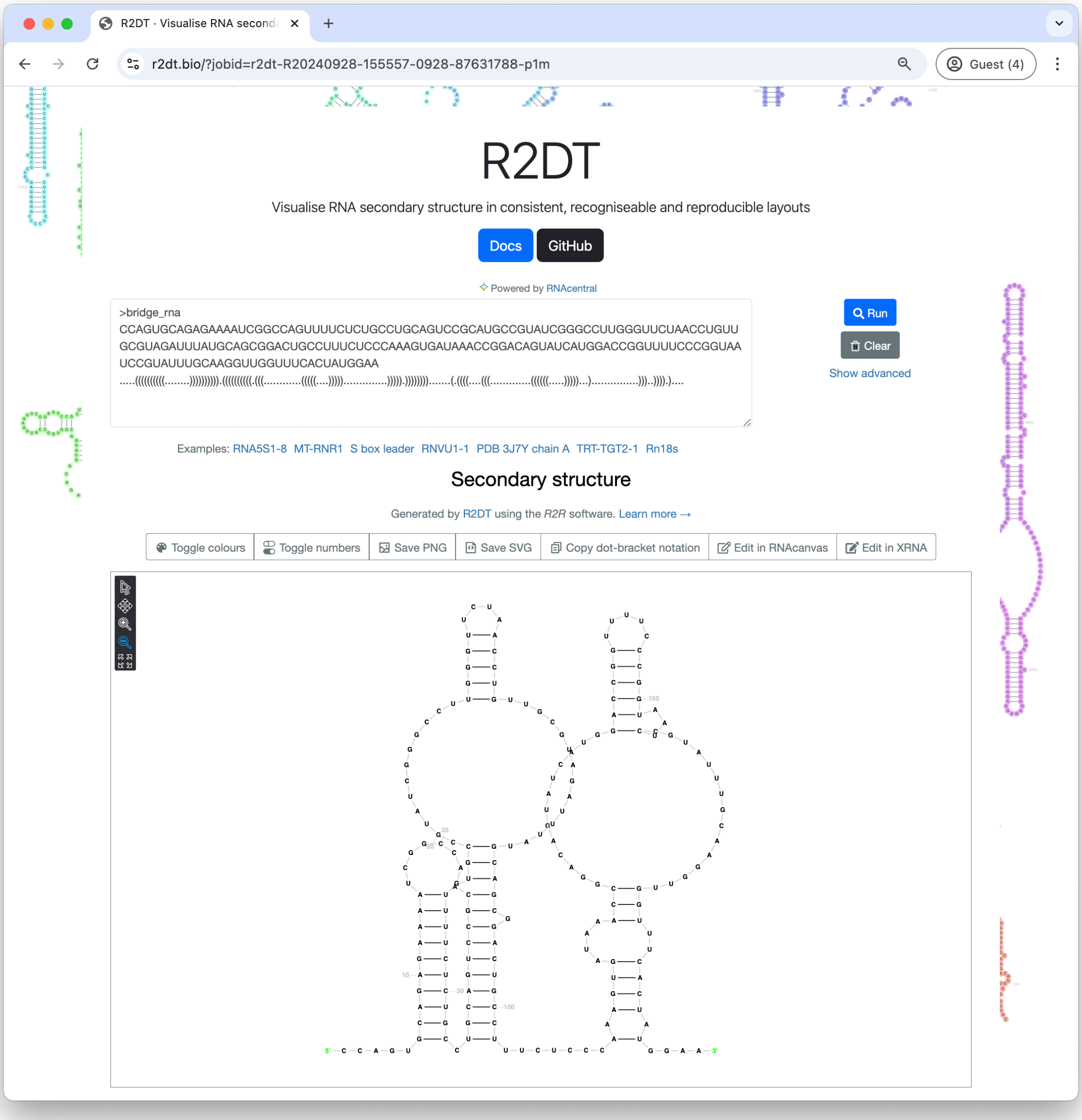


In the following steps, we will edit this R2DT results to generate a new 2D diagram that avoids structural overlaps.

**STEP 2.** Click the **Edit in RNAcanvas** button and copy the `rna_2d_schema_url` parameter from the URL.

Example URL: `https://www.ebi.ac.uk/Tools/services/rest/r2dt/result/r2dt-R20240928-155557-0928-87631788-p1m/json`

Note that the R2DT results expire after 7 days so the instructions work only with recently generated R2DT API URLs.

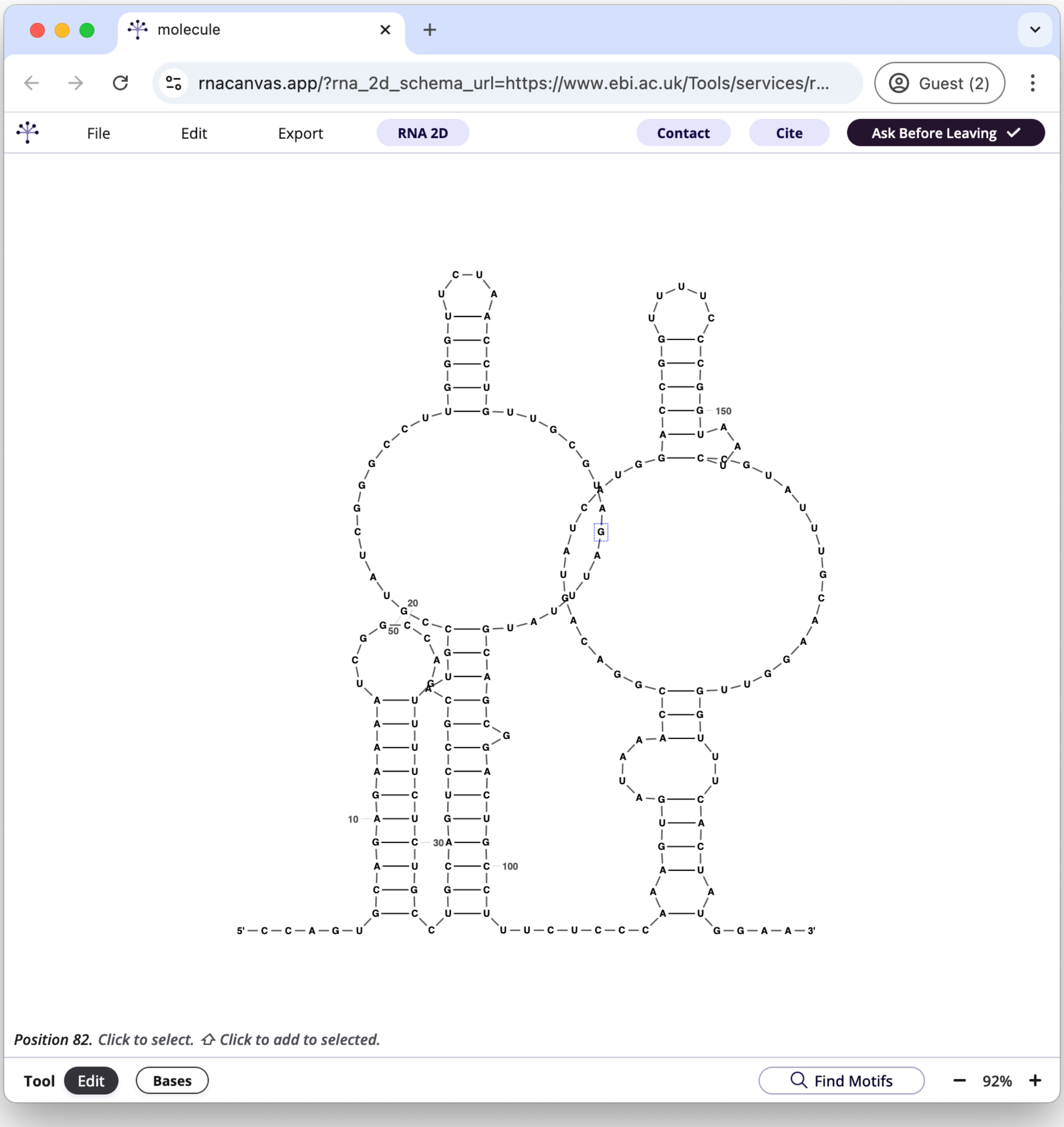

**STEP 3.** Navigate to RNAcanvas Code at https://code.rnacanvas.app, open JavaScript Console and type `assistant()`:

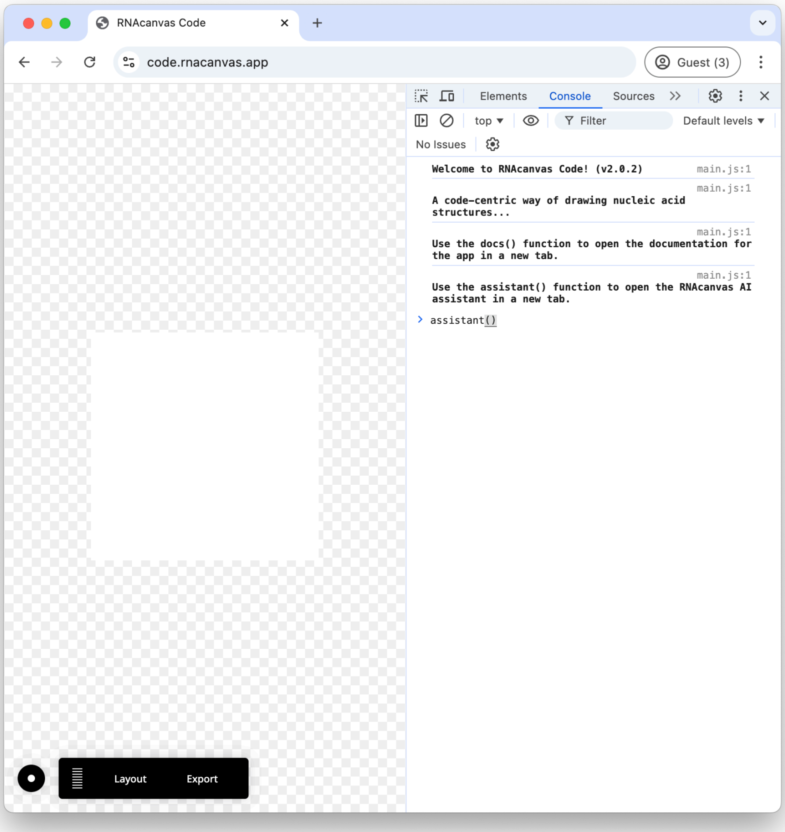


This will launch the RNAcanvas ChatGPT interface in a new browser window. Authenticate with a free OpenAI user account to get access to the prompt interface:

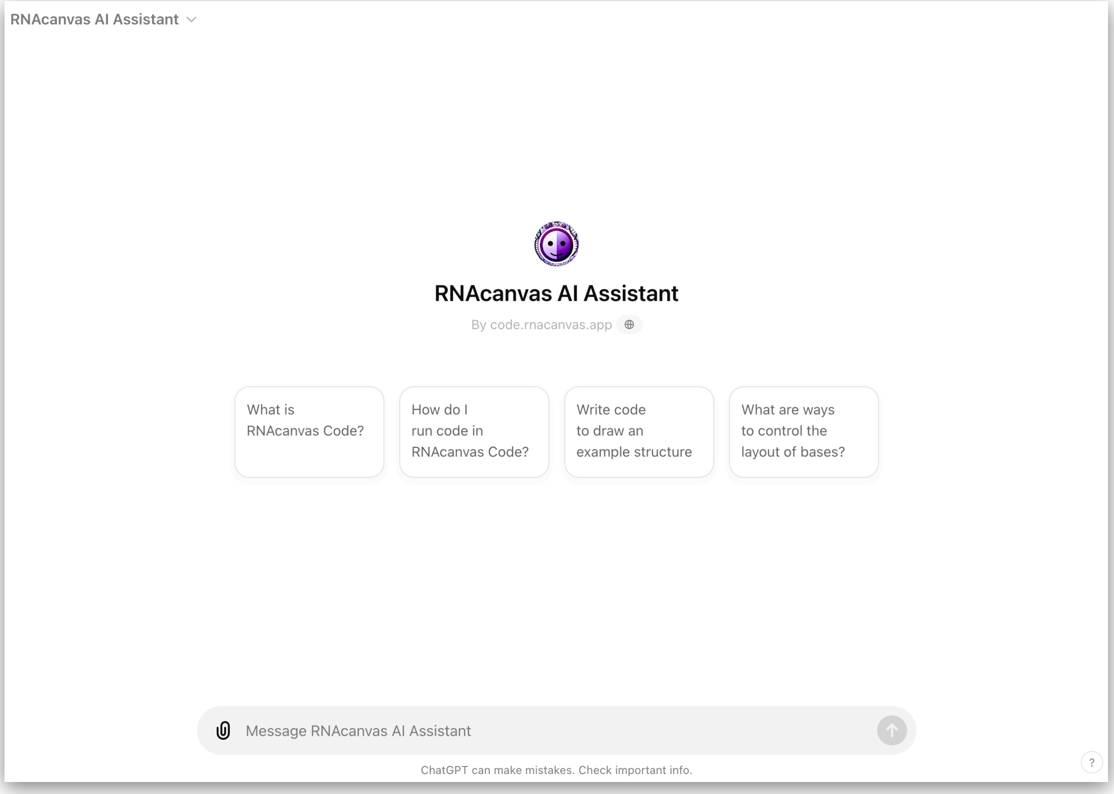

**STEP 4.** Use ChatGPT to load R2DT diagram into RNAcanvas Code using the URL obtained at step 2.

> Load RNA structure using the following RNA 2D JSON Schema file: https://www.ebi.ac.uk/Tools/services/rest/r2dt/result/r2dt-R20240928-155557-0928-87631788-p1m/json

To load an RNA structure using the RNA 2D JSON Schema from the provided URL, you can use the `fetch` method along with the RNAcanvas Code API. Here's the JavaScript code you can run in the web browser console to load the RNA structure:

javascript    Copy code

```javascript
// the URL of the RNA 2D JSON schema
var schemaURL = 'https://www.ebi.ac.uk/Tools/services/rest/r2dt/result/r2dt-R20240

fetch(schemaURL)
  .then(response => response.text())
  .then(text => app.drawSchema(JSON.parse(text)))
  // ensure the drawing is big enough to fit the structure
  .then(() => app.drawing.setPadding(1000))
  // fit the view of the drawing to the structure
  .then(() => app.drawingView.fitToContent());
```

This code will fetch the RNA 2D JSON from the URL, parse it, and render the structure using the RNAcanvas API. You can adjust the padding as needed to fit the structure into the visible drawing area.

Copy the JavaScript code from ChatGPT and paste into RNAcanvas Code:

```javascript
var schemaURL = 'https://www.ebi.ac.uk/Tools/services/rest/r2dt/result/r2dt-R20240928-155557-0928-87631788-p1m/json';

fetch(schemaURL)
  .then(response => response.text())
  .then(text => app.drawSchema(JSON.parse(text)))
  // ensure the drawing is big enough to fit the structure
  .then(() => app.drawing.setPadding(1000))
  // fit the view of the drawing to the structure
  .then(() => app.drawingView.fitToContent());
```

The diagram has been loaded and is ready to be edited to remove the overlaps:

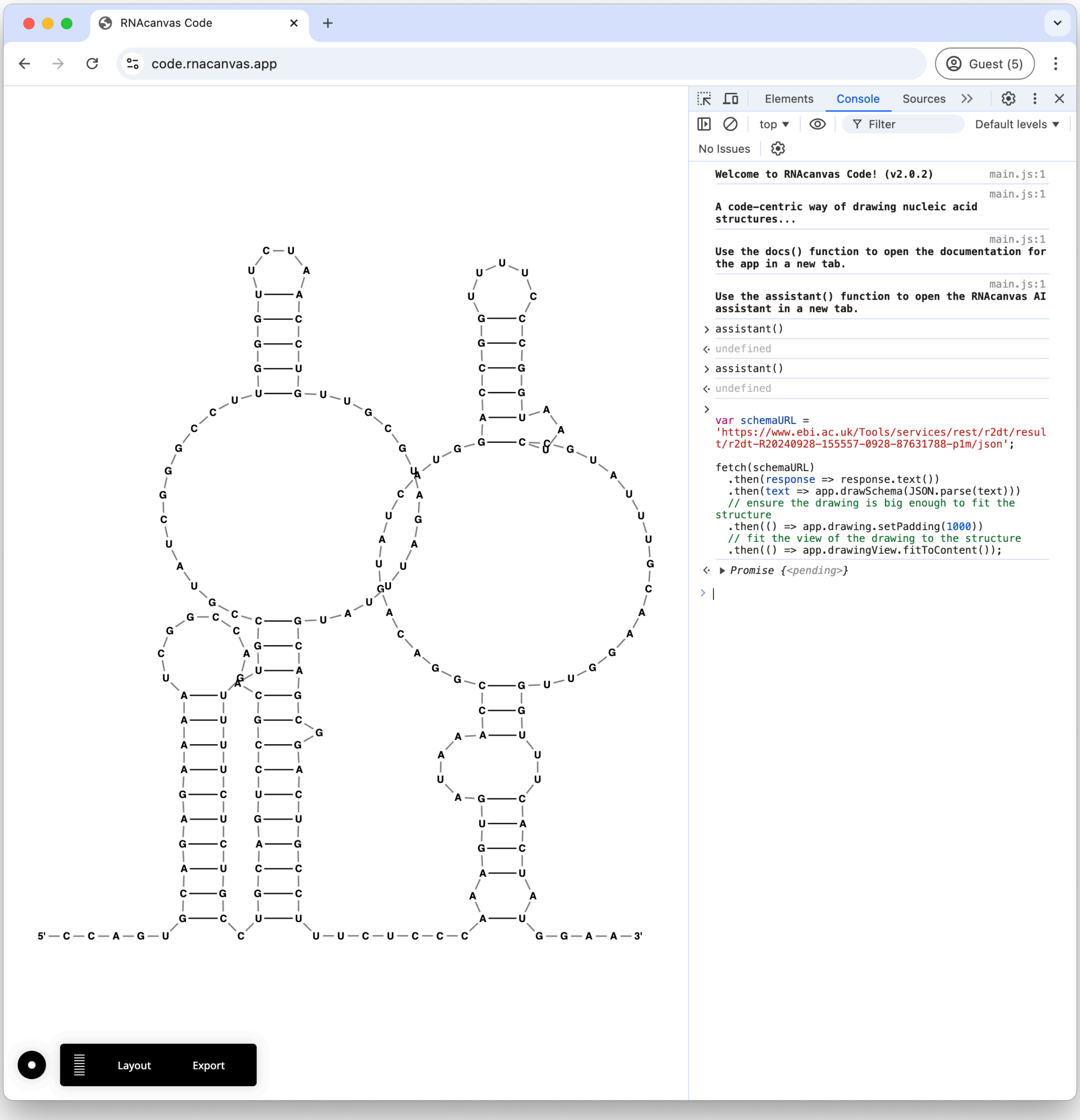

**STEP 5**. Move the first 33 nucleotides to the left by 10px. Nucleotide numbers can be obtained by hovering the mouse over nucleotides in the R2DT interface from Step 1.

move first 33 nucleotides to the left by 10px

Here is the JavaScript code to move the first 33 nucleotides to the left by 10px in the current RNA drawing:

javascript Copy code

```javascript
// Get all the bases in the drawing
var bases = [...app.drawing.bases];

// Move the first 33 nucleotides 10px to the left
bases.slice(0, 33).forEach(base => {
  let center = base.getCenterPoint();
  base.setCenterPoint({ x: center.x - 10, y: center.y });
});
```

#### Explanation:

- The `slice(0, 33)` method selects the first 33 nucleotides.
- The `getCenterPoint()` method retrieves the current position of each base.
- The `setCenterPoint()` method shifts the X-coordinate of each base by 10px to the left (subtracting 10 from the X value).

You can run this code in the web browser console to apply the movement to the first 33 nucleotides.

Code generated by ChatGPT:

```javascript
// Get all the bases in the drawing
var bases = [...app.drawing.bases];

// Move the first 33 nucleotides 10px to the left
bases.slice(0, 33).forEach(base => {
  let center = base.getCenterPoint();
  base.setCenterPoint({ x: center.x - 10, y: center.y });
});
```

### Resulting diagram:

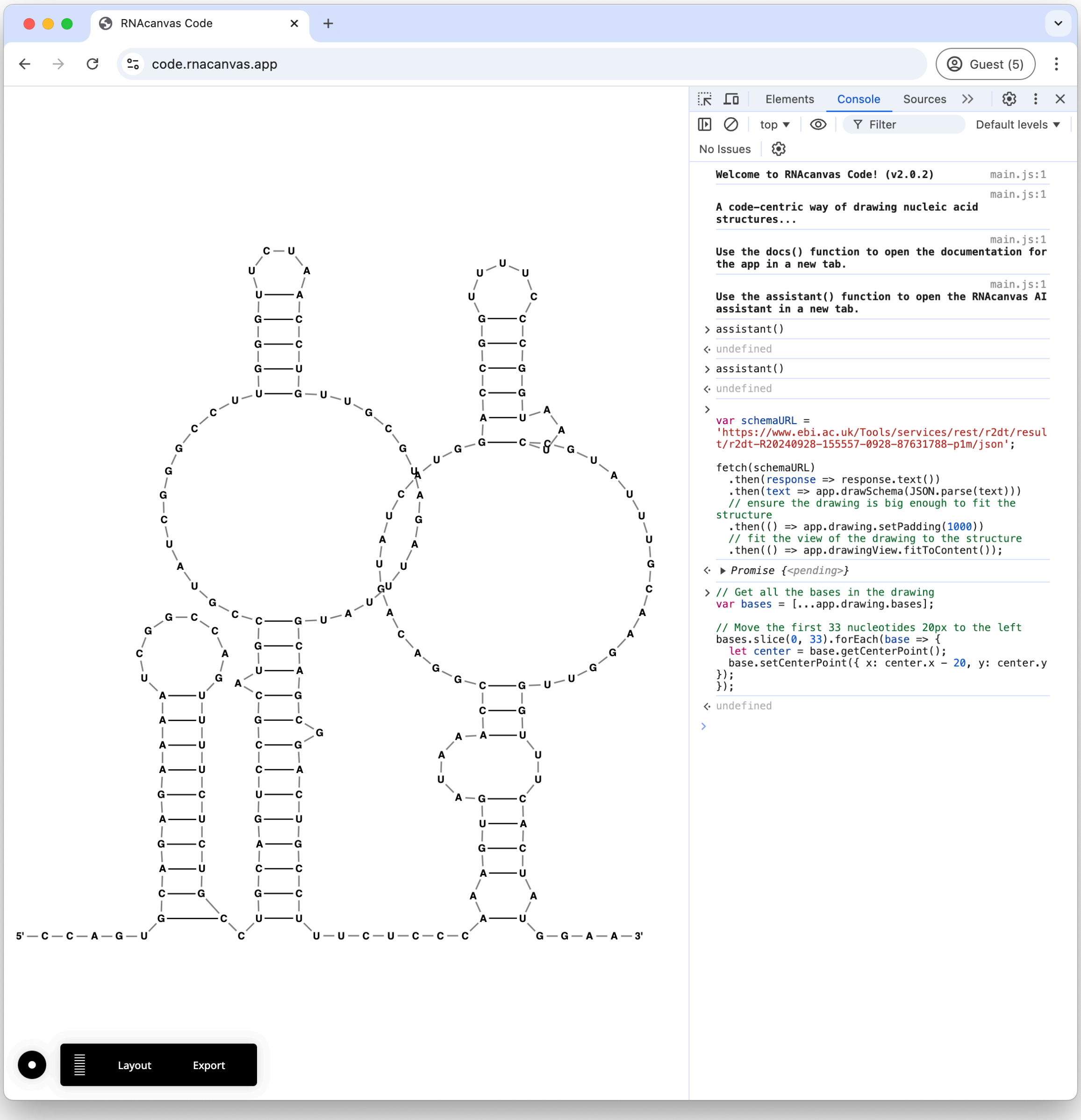

### STEP 6. Straighten nucleotides 123 to 135.

straighten nucleotides 123 to 135

Here is the JavaScript code to straighten nucleotides 123 to 135 in the current RNA drawing:

javascript    Copy code

```javascript
// Get all the bases in the drawing
var bases = [...app.drawing.bases];

// Select nucleotides 123 to 135 (note: 0-based index, so adjust accordingly)
var selectedBases = bases.slice(122, 135);  // Nucleotides 123 to 135

// Straighten the selected bases by evenly spacing them
let firstBaseCenter = selectedBases[0].getCenterPoint();
let lastBaseCenter = selectedBases[selectedBases.length - 1].getCenterPoint();

// Calculate the spacing between each base
let spacingX = (lastBaseCenter.x - firstBaseCenter.x) / (selectedBases.length - 1);
let spacingY = (lastBaseCenter.y - firstBaseCenter.y) / (selectedBases.length - 1);

// Set the position of each base along a straight line
selectedBases.forEach((base, i) => {
  base.setCenterPoint({
    x: firstBaseCenter.x + i * spacingX,
    y: firstBaseCenter.y + i * spacingY,
  });
});
```

#### Explanation:

- `slice(122, 135)` selects the nucleotides from position 123 to 135 (adjusting for 0-based indexing).
- The script calculates the spacing between the first and last selected nucleotides, dividing it evenly for all bases.
- Each base's position is updated to create a straight line between the first and last bases.

Code generated by ChatGPT:

```javascript
// Get all the bases in the drawing
var bases = [...app.drawing.bases];

// Select nucleotides 123 to 135 (note: 0-based index, so adjust accordingly)
var selectedBases = bases.slice(122, 137);  // Nucleotides 123 to 135

// Straighten the selected bases by evenly spacing them
let firstBaseCenter = selectedBases[0].getCenterPoint();
let lastBaseCenter = selectedBases[selectedBases.length - 1].getCenterPoint();

// Calculate the spacing between each base
let spacingX = (lastBaseCenter.x - firstBaseCenter.x) / (selectedBases.length - 1);
let spacingY = (lastBaseCenter.y - firstBaseCenter.y) / (selectedBases.length - 1);

// Set the position of each base along a straight line
selectedBases.forEach((base, i) => {
  base.setCenterPoint({
    x: firstBaseCenter.x + i * spacingX,
    y: firstBaseCenter.y + i * spacingY,
  });
});
```

**Resulting diagram:**

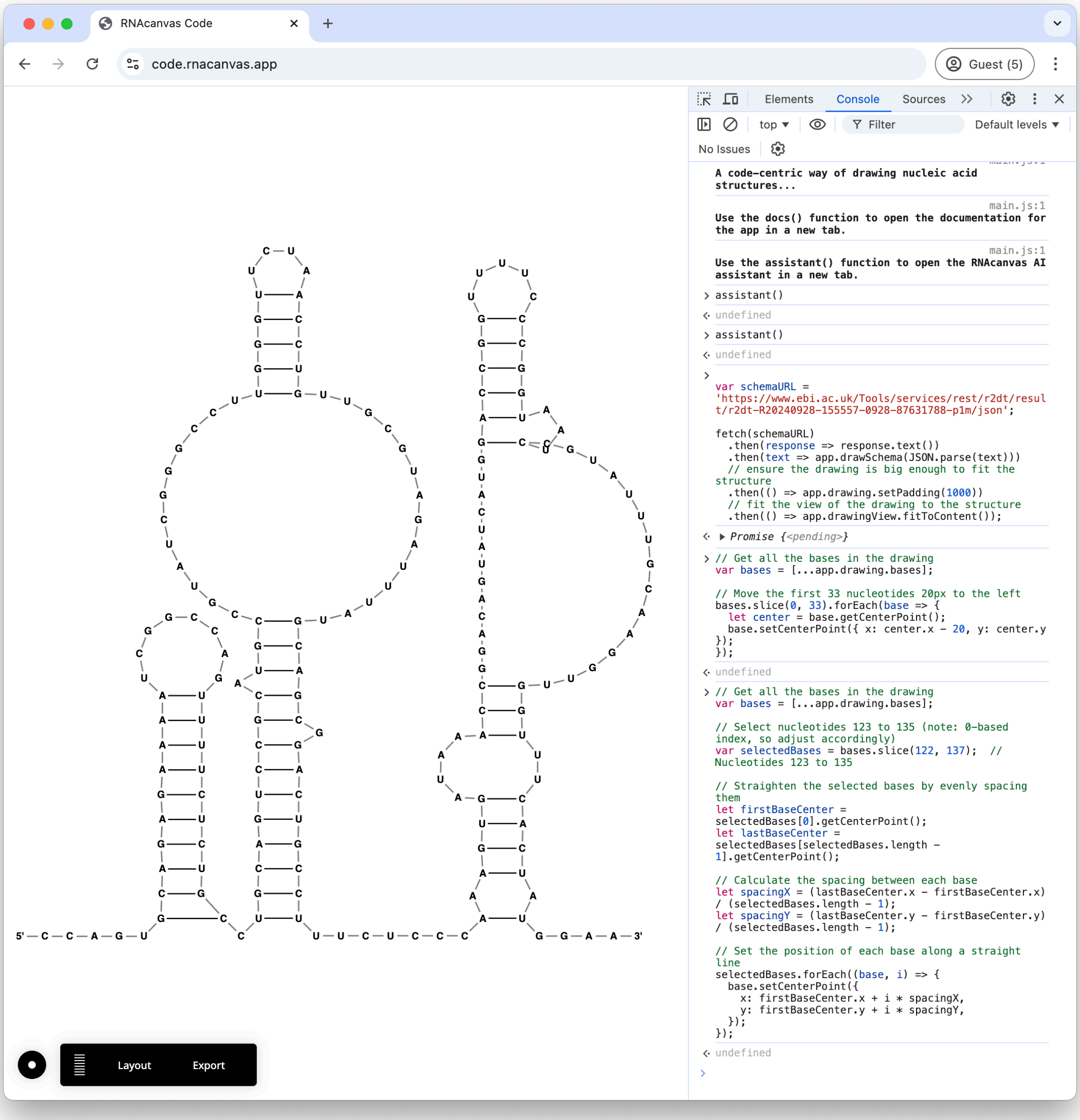


The diagram no longer has major overlaps and can be further edited manually or by using ChatGPT.

### CONCLUSIONS

This example highlights the advantages of using natural language for editing RNA 2D diagrams. Users can manipulate structures by describing the desired outcome, eliminating the need to manually select nucleotides, navigate through user interface controls, and apply operations.

As LLM technology and available tools continue to evolve, the process will be further streamlined, enabling direct diagram editing without requiring users to interact with the underlying code.
